# DeePVP: Identification and classification of phage virion protein using deep learning

**DOI:** 10.1101/2021.10.23.465539

**Authors:** Zhencheng Fang, Tao Feng, Hongwei Zhou

## Abstract

The poor annotation of phage virion protein (PVP) is the bottleneck of many areas of viral research, such as viral phylogenetic analysis, viral host identification and antibacterial drug design. Because of the high diversity of the PVP sequences, the PVP annotation remains a great challenging bioinformatic task. Based on deep learning, we present DeePVP that contains a main module and an extended module. The main module aims to identify the PVPs from non-PVP over a phage genome, while the extended module can further classify the predicted PVP into one of the ten major classes of PVP. Compared with the state-of-the-art tools that can distinguish PVP from non-PVP, DeePVP’s main module performs much better, with an *F1-score* 9.05% higher in the PVP identification task. Compared with PhANNs, a tool that can further classify the predicted PVP into a specific class, the overall accuracy of DeePVP’s extended module is approximately 3.72% higher in the PVP classification task. Two application cases on the genome of mycobacteriophage PDRPxv and *Escherichia* phage HP3 show that the predictions of DeePVP are much more reliable and can better reveal the compact PVP-enriched region, which may be conserved during the viral evolution process, over the phage genome.

## INTRODUCTION

Viruses are the most dominant biological entities in the biosphere (1). Because of the diversity of phage genomes, there are about 50–90% of phage genes cannot be assigned function (2). Additionally, it has been estimated that approximately 60%-99% of viral metagenomic data do not have obvious homology regions to the known database (3). These results mean that a large number of viral genes exist as “dark matter”, which is a barrier to our understanding of viral genomes. Therefore, the development of gene function prediction tools for the viral genome is urgently needed.

Phage virion proteins (PVPs), also called phage structural proteins, refer to the proteins that make up of viral particles such as the head and tail. The comprehensive annotation of PVPs is essential for many phage genome analyses (4). For example, phages lack marker genes such as 16S rDNA in bacteria, and it has been suggested that some PVPs may be used as marker genes for phage genome analysis (5). Additionally, PVPs in the phage genome could help to improve our understanding of phage-bacterial host interactions (6), direct antibacterial drug and antibiotic design (7), select specific phages for phage therapy (8) and assist in prophage identification in the bacterial genome (9).

In the past, the experimental method of mass spectrometry has been commonly used in the task of PVP identification (10). With the rapid increase in viral sequencing data, low-cost and high-performing bioinformatic algorithms to perform PVP annotation are urgently needed in this field. However, the diversity of PVPs is much higher than that of the enzymes encoded in the phage genome, which makes the identification of PVPs much more difficult (11). To overcome the difficulty of PVP identification, several de novo algorithms for PVP identification have been proposed (8,11-23). Most of these tools are two-class classifiers to distinguish whether the given phage protein is PVP. These tools primarily first constructed a benchmark training and testing dataset containing PVP and non-PVP from the public database and used specific machine learning-based algorithms, such as support vector machine (SVM), to train and test the classifier using the dataset. Among these tools mentioned above, PVPred (13), PVP-SVM (15), PVPred-SCM (20), Meta-iPVP (21) and VirionFinder (22) are available via a one-click software package or a web server during the period of this work. In addition to these two class classification tools that distinguish PVP and non-PVP, some tools that are designed for identifying specific PVPs, such as the capsid and tail, have also been developed (5,11). Different from these tools, the PhANNs tool (8) is a multiclass classifier that can not only identify whether a given protein is PVP but also further classify the predicted PVP into several major PVP classes, making PhANNs a more powerful tool for PVP annotation.

Although these tools can achieve better performance during the period of their publication, further effort can be made to improve the PVP annotation. For example, most of these tools were trained and tested using a small-scale dataset containing less than 1000 proteins, and such a dataset may not reflect the diversity of the PVP sequence features. With the rapid growth of the public database size, more PVPs and non-PVPs can be included to develop the algorithm. In terms of the characterization method for the protein sequence, most tools used amino acid composition-based vectors, such as the *k*-mer vector used by PhANNs, to characterize the sequence. However, such a vector may be sparse, which is difficult to fit by the algorithm. Many current tools mentioned above have to perform the feature selection step before the feature vector is imported to the algorithm. Moreover, except for PhANNs, current PVP annotation tools can only distinguish PVP and non-PVP or identify a certain class of PVP. They cannot further classify PVP into specific classes, which prevents detailed analysis of the phage genome.

To improve the performance of PVP annotation, we present DeePVP. DeePVP takes a phage protein as input. For each input protein, the main module of DeePVP outputs a PVP likelihood score to reflect whether the given protein belongs to the PVP, while the extended module further calculates whether a predicted PVP belongs to one of the following ten major classes of PVP, namely, head-tail joining, collar, tail sheath, tail fiber, portal, minor tail, major tail, baseplate, minor capsid, and major capsid, which are the dominant categories in PVP. Both modules use the one-hot encoding form to characterize the protein sequence and use a convolution neural network (CNN) as the classifier to extract the abstract feature of the protein. Testing using the benchmark dataset and two application cases demonstrates the advantage of DeePVP in PVP annotation. DeePVP is freely available at https://github.com/zhenchengfang/DeePVP/.

## MATERIAL AND METHODS

Several benchmark training and testing datasets of PVP and non-PVP have been constructed in previous work (8,11-23). In this work, the benchmark dataset of PhANNs (http://edwards.sdsu.edu/phanns/download/expandedDB.tgz) (8), which was constructed using proteins from the NCBI protein database and GenBank database, was used to develop DeePVP. This dataset is chosen for the following reasons: (1) during the period of this work, the PhANNs dataset is the largest PVP and non-PVP dataset, containing a total of 168,660 PVP and 369,553 non-PVP, while most of the other datasets contain less than 1000 protein; (2) the homology between the training and testing proteins is less than 40%, which is important to evaluate whether the algorithm can predict the novel proteins; and (3) the dataset includes all 10 major classes of PVPs that we focused on.

The framework of DeePVP is shown in Figure 1. Selecting a proper representation method for biological sequences is an important step for algorithm development. Although the *k*-mer frequency vector has been widely used in many studies, such global statistics may lose some local information of the sequence, such as the information of some conserved domains or motifs of the sequence (24). In DeePVP, we used the “one-hot” encoding form to represent the protein sequence. In this way, each amino acid is represented by a “one-hot” vector containing 20 bits, in which 19 bits are 0 and a certain bit is 1; therefore, the information of each amino acid is retained in the digitized model (see Supplementary Data S1 for more details). The “one-hot” encoded protein will first go through the main module. The main module uses CNN to extract the sequence features to just whether the given protein is PVP. The CNN contains a 1D convolution layer, 1D global max pooling layer, batch normalization layer and full connection layer, and finally, the sigmoid layer outputs a PVP score between 0-1. By default, a protein with a PVP score higher than 0.5 is regarded as PVP. The extended module also uses CNN to classify the protein into a specific class of PVP. In the extended module, the CNN contains a 1D convolution layer, 1D global max pooling layer, batch normalization layer and full connection layer, and finally, the softmax layer outputs 10 likelihood scores representing the probability of the protein belonging to the head-tail joining, collar, tail sheath, tail fiber, portal, minor tail, major tail, baseplate, minor capsid, and major capsid. By default, the category with the highest score will be chosen as the final prediction. The details of the hyperparameter selection are provided in Supplementary Data S1.

**Figure 1.**
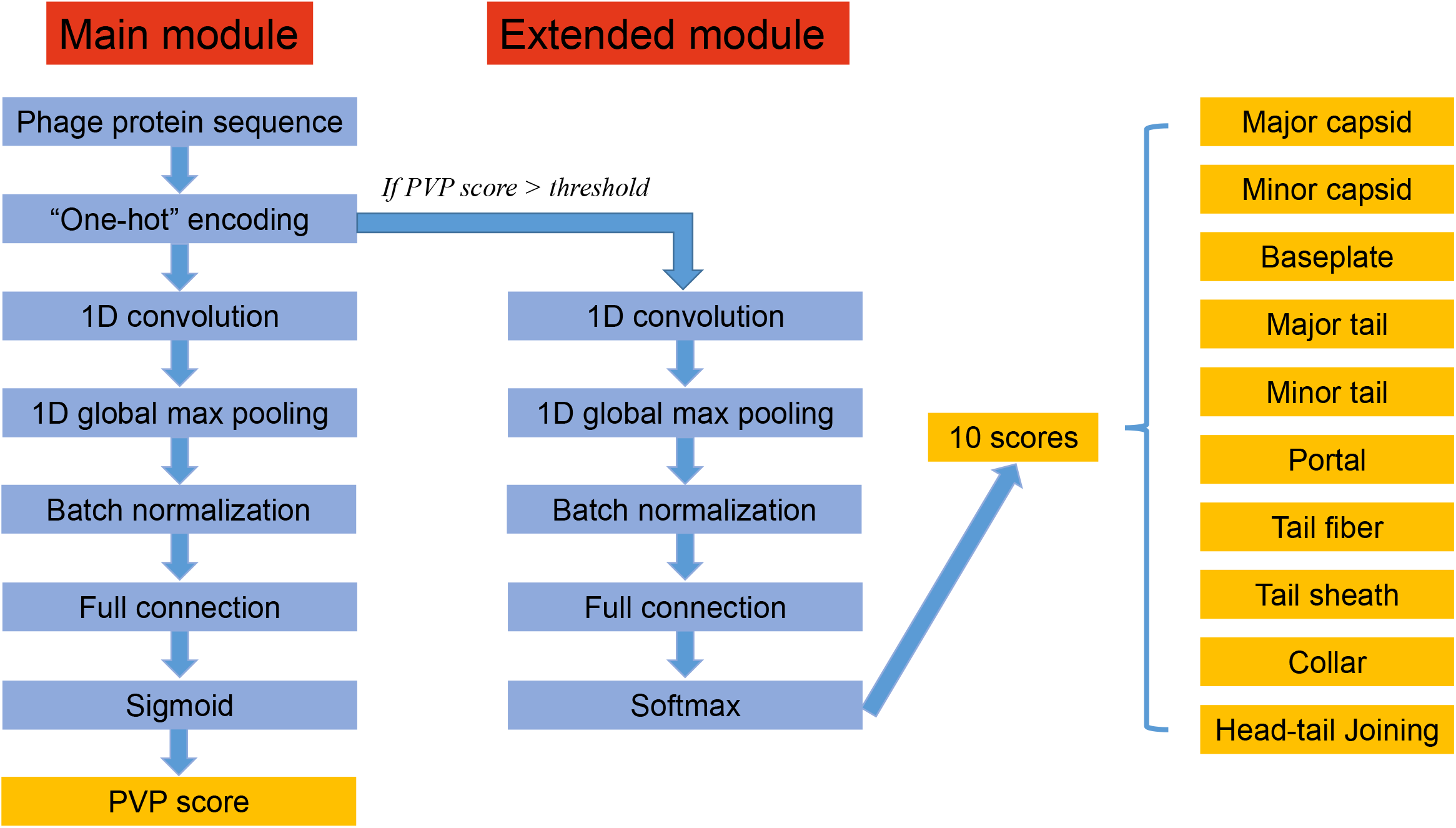
The frame work of DeePVP. DeePVP takes a protein sequence as input, the main module calculates a PVP score representing the probability that the protein belongs to PVP, and the extended module calculates 10 likelihood scores representing which category the predicted PVP belongs to.

In the training process, both PVPs and non-PVPs in the training set were used to train the main module, and only PVPs were used to train the extended module. In the prediction process, each protein will go through the main module and extended module. However, it is worth noting that if a protein obtains a PVP score lower than the threshold, the 10 scores calculated by the extended module do not make sense because this protein does not belong to PVP. Therefore, in the workflow of DeePVP, we will normalize the 10 scores from the extended module to make them sum up to the PVP score.

## RESULTS

### Performance comparison in the PVP identification task

Considering that most of the current tools can only identify PVPs from non-PVPs, we first compared the performance between DeePVP’s main module and the related available tools, namely, PVPred, PVP-SVM, PVPred-SCM, Meta-iPVP, PhANNs and VirionFinder, using the test set. The evaluation criteria are:

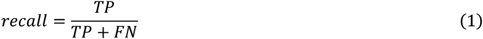

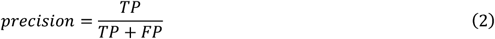

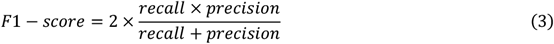

where *TP, TN, FN* and *FP* represent the number of true positive, true negative, false negative and false positive predictions, respectively. Among these three criteria, the *F1-score* can serve as the comprehensive index to evaluate the tools. The performance comparison is shown in the Table 1. Although the *recall* of DeePVP is slightly lower than that of PhANNs and VirionFinder, the *precision* of DeePVP is much better than all the other tools, and the comprehensive index of *F1-score* is 9.05% much higher than that of the tool PhANNs, which performs the best among the comparative tools. Since DeePVP and PhANNs were trained using the same training data, the advantage of DeePVP showed that the “one-hot” encoding form may be a more detailed representation method than the *k*-mer frequency used in PhANNs to characterize the protein sequence, and the deep convolution neural network in DeePVP may be more powerful than the simple shallow network containing 2 hidden layers used in PhANNs for feature extraction.

**Table 1.** Performance comparison of DeePVP and related tools in the PVP identification task.

| Tool | *Recall (%)* | *Precision (%)* | *F1-score (%)* |
| --- | --- | --- | --- |
| **DeePVP** | **88.10** | **96.75** | **92.22** |
| VirionFinder | 90.91 | 44.00 | 59.30 |
| PhANNs | 91.68 | 76.11 | 83.17 |
| Meta-iPVP | 82.41 | 53.29 | 64.72 |
| PVPred-SCM | 41.28 | 38.71 | 39.95 |
| PVP-SVM | 41.31 | 46.19 | 43.62 |
| PVPred | 29.84 | 42.23 | 34.97 |

### Performance comparison in the PVP classification task

We further compared the performance between DeePVP’s extended module and PhANNs in the PVP classification task. Since all the other tools cannot further classify a given PVP into a specific class, these tools are not included in this analysis. During the comparison, we assume that all the PVPs have already been correctly predicted in the upstream analysis; therefore, non-PVPs are excluded in the test set in this subsection. Actually, in the released package of DeePVP, the main module and the extended module can be run using an integrated pipeline or be run separately. For example, if researchers have already identified PVPs using related computational or experimental methods, such as mass spectrometry, they can directly run DeePVP’s extended module for PVP classification.

We used the criteria of *recall, precision* and *F1-score* to evaluate the performance of DeePVP and PhANNs in each PVP category. We also used the *accuracy* to evaluate the overall performance of the tools. The *accuracy* is defined as the ratio of the number of correctly classified PVPs to the total PVPs. The performance comparison is shown in the Table 2. The overall *accuracy* of DeePVP is 3.72% higher than that of PhANNs, indicating that DeePVP has a better ability to classify PVP. In terms of each category, although the *F1-score* of DeePVP in the major tail, tail fiber, tail sheath and collar is lower than that of PhANNs, DeePVP presents a higher *F1-score* in other categories. Although the advantage of DeePVP’s *accuracy* is not quite obvious, with only 3.72% higher than PhANNs, it is worth noting that such performance is evaluated under the assumption that all PVPs have been correctly predicted. Since DeePVP performs much better than PhANNs in the PVP identification task, we consider that the comprehensive performance of PVP annotation of DeePVP is better than PhANNs.

**Table 2.** Performance comparison of DeePVP and PhANNs in the PVP classification task.

| Category | Tool | *Recall (%)* | *Precision (%)* | *F1-score (%)* | *Accuracy (%)* |
| --- | --- | --- | --- | --- | --- |
| Major capsid | DeePVP | 98.58 | 90.88 | 94.57 | NA |
|  | PhANNs | 91.98 | 94.24 | 93.10 |  |
| Minor capsid | DeePVP | 50.62 | 44.09 | 47.13 |  |
|  | PhANNs | 80.25 | 13.98 | 23.81 |  |
| Baseplate | DeePVP | 86.49 | 95.96 | 90.98 |  |
|  | PhANNs | 78.85 | 88.76 | 83.51 |  |
| Major tail | DeePVP | 77.09 | 61.23 | 68.25 |  |
|  | PhANNs | 80.48 | 86.88 | 83.56 |  |
| Minor tail | DeePVP | 90.28 | 93.24 | 91.74 |  |
|  | PhANNs | 85.42 | 90.50 | 87.89 |  |
| Portal | DeePVP | 93.10 | 93.62 | 93.36 |  |
|  | PhANNs | 85.88 | 93.77 | 89.65 |  |
| Tail fiber | DeePVP | 78.24 | 68.42 | 73.00 |  |
|  | PhANNs | 77.93 | 69.66 | 73.56 |  |
| Tail sheath | DeePVP | 92.86 | 99.42 | 96.03 |  |
|  | PhANNs | 94.39 | 99.64 | 96.94 |  |
| Collar | DeePVP | 36.33 | 80.15 | 50.00 |  |
|  | PhANNs | 87.00 | 75.65 | 80.93 |  |
| Head-Tail Joining | DeePVP | 96.48 | 96.33 | 96.40 |  |
|  | PhANNs | 88.49 | 71.88 | 79.33 |  |
| All | DeePVP |  | NA |  | **91.06** |
|  | PhANNs |  |  |  | 87.34 |

### Application case 1: PVP annotation of the mycobacteriophage PDRPxv genome

To demonstrate the value of DeePVP in PVP annotation and its reliability, we first used DeePVP and related tools to perform PVP annotation on the mycobacteriophage PDRPxv genome (GenBank accession: KR029087), a phage that may be a candidate therapeutic for pathogenic *Mycobacterium* species (25). In the PDRPxv genome, PVPs have been identified by the experimental method of mass spectrometry; therefore, such experimental data can be used to evaluate the reliability of computational tools. The tools DeePVP, PVPred, PVP-SVM, PVPred-SCM, Meta-iPVP, PhANNs and VirionFinder were used to perform PVP annotation on the PDRPxv genome, and the mass spectrometry data were used to evaluate the *recall, precision* and *F1-score* of each tool. The performance of each tool in the PVP identification task is shown in Table 3. We found that most of the comparative tools did not perform well, with *F1-scores* lower than 50%. PhANNs performed better than the other comparative tools, and the *F1-score* of DeePVP was 21.19% much higher than that of PhANNs, indicating that DeePVP provides a more reliable prediction.

**Table 3.**
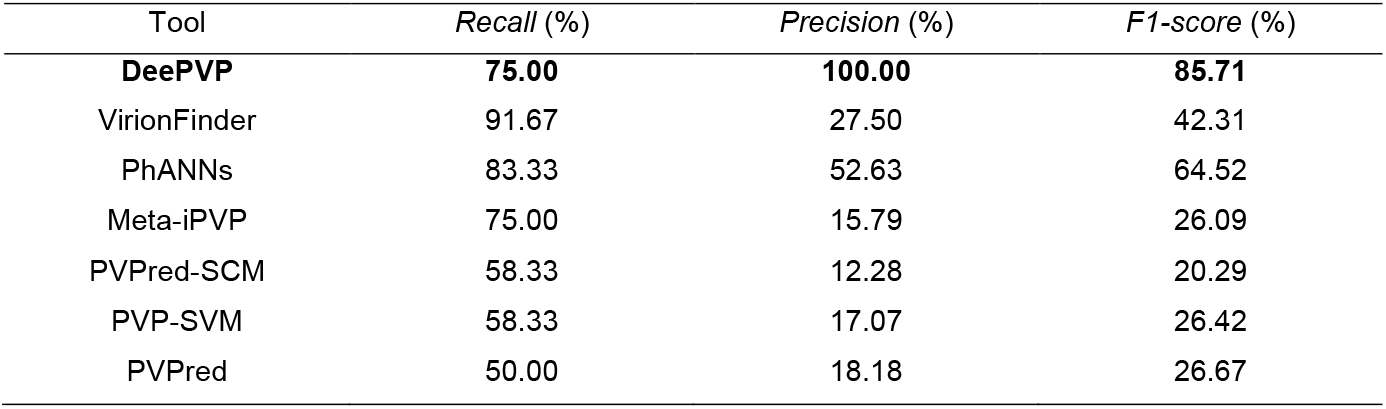
Performance comparison in the PVP identification task on the PDRPxv genome based on mass spectrometry data.

In Figure 2, we show the base coordinates for the PVPs uncovered by mass spectrometry and the predicted PVPs of each tool. The mass spectrometry data showed that all PVPs were located within a compact PVP-enriched region, and outside the PVP-enriched region, there was no PVP. In fact, it has been shown that many PVPs often encode next to each other over the genome (26,27), and such PVP distribution patterns may be general in other phages. Interestingly, we found that the PVP distribution pattern revealed by DeePVP was quite consistent with that revealed by mass spectrometry. Although DeePVP failed to predict a few PVPs, all the other predicted PVPs were located within the PVP-enriched region. In contrast, the PVPs predicted by the other tools seem to be distributed randomly on the genome. This phenomenon shows that the prediction of DeePVP is more convincing and that DeePVP has a better ability to reveal the genomic structure of a phage genome.

**Figure 2.**
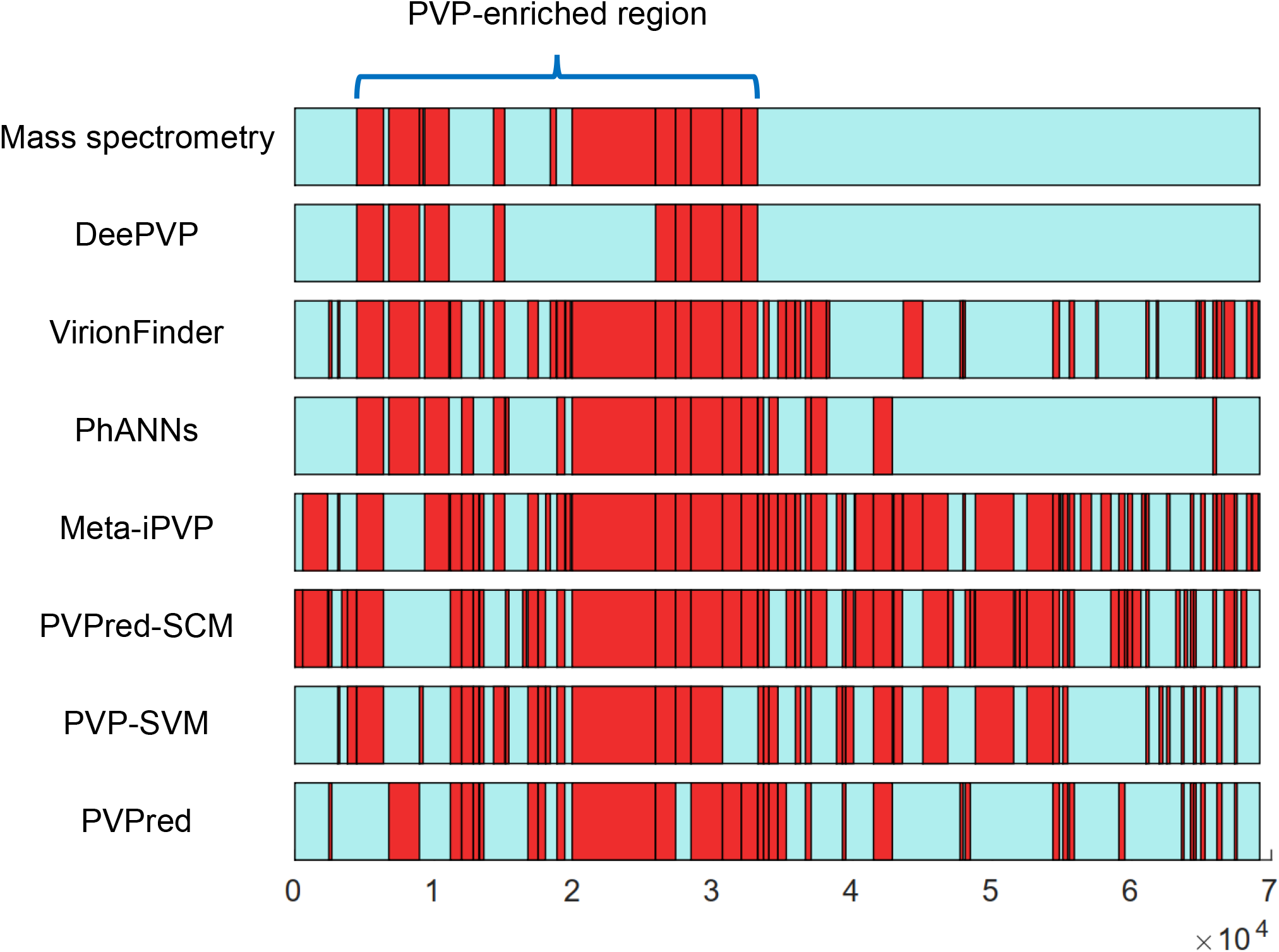
The distribution of the PVPs uncovered by mass spectrometry and the PVPs predicted by each tool on the mycobacteriophage PDRPxv genome. The blue bars represent the PDRPxv genome, and the red boxes represent the PVPs. The horizontal axis represents the base coordinates.

According to mass spectrometry, the PDRPxv genome contains 12 PVPs. Sinha *et al*. (25) inferred the putative function of each PVP by combining several strategies, including domain search, homology analysis, adjacent gene analysis and protein secondary structure analysis. In Table 4, we compared the PVP class predicted by DeePVP and PhANNs with the putative function revealed by Sinha *et al*. We found that the predictions of DeePVP and PhANNs were basically consistent with the putative function. In particular, 5 putative minor tail proteins are encoded continuously (Gp29-Gp33) in the genome, and both DeePVP and PhANNs could predict this minor tail protein cluster appropriately.

**Table 4.**
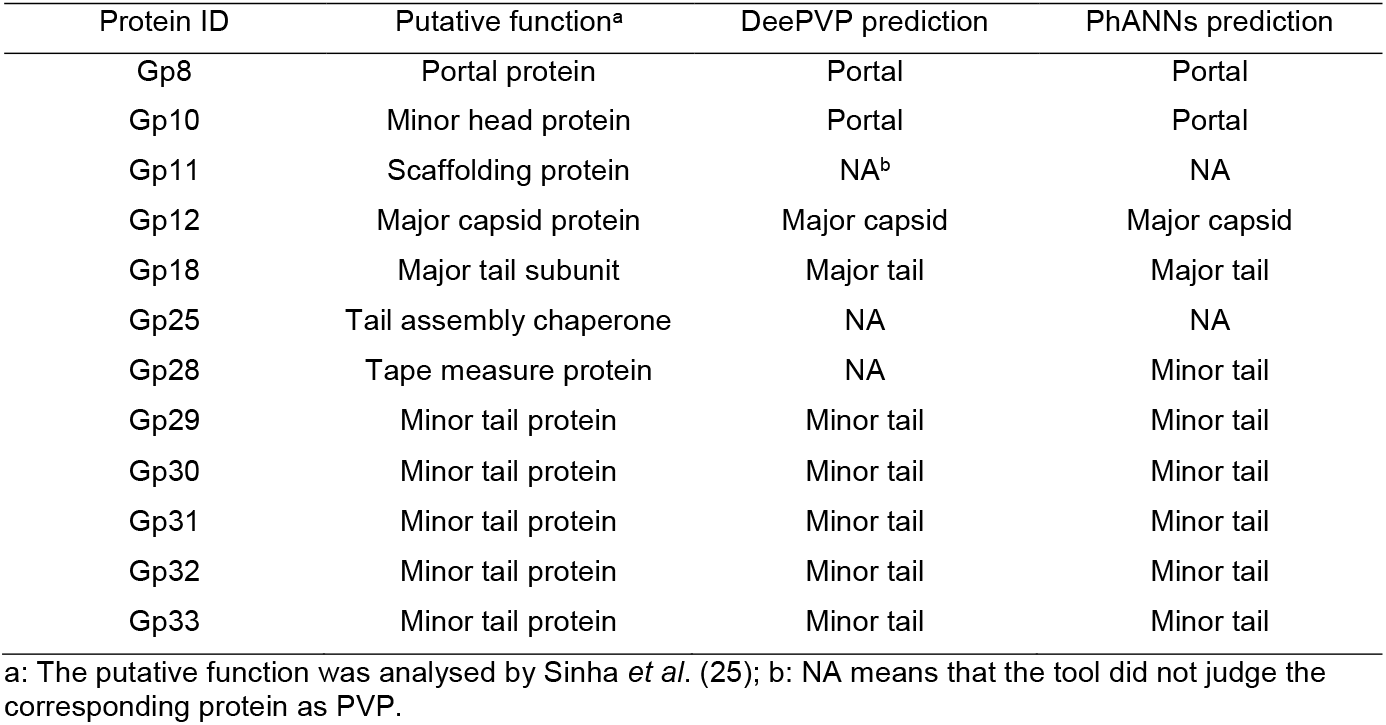
PDRPxv genome PVP classification by DeePVP and PhANNs.

| Protein ID | Putative function<sup>a</sup> | DeePVP prediction | PhANNs prediction |
| --- | --- | --- | --- |
| Gp8 | Portal protein | Portal | Portal |
| Gp10 | Minor head protein | Portal | Portal |
| Gp11 | Scaffolding protein | NA<sup>b</sup> | NA |
| Gp12 | Major capsid protein | Major capsid | Major capsid |
| Gp18 | Major tail subunit | Major tail | Major tail |
| Gp25 | Tail assembly chaperone | NA | NA |
| Gp28 | Tape measure protein | NA | Minor tail |
| Gp29 | Minor tail protein | Minor tail | Minor tail |
| Gp30 | Minor tail protein | Minor tail | Minor tail |
| Gp31 | Minor tail protein | Minor tail | Minor tail |
| Gp32 | Minor tail protein | Minor tail | Minor tail |
| Gp33 | Minor tail protein | Minor tail | Minor tail |

a: The putative function was analysed by Sinha *et al.* (25); b: NA means that the tool did not judge the corresponding protein as PVP.

### Application case 2: A novel conserved PVP-enriched region was found in *Escherichia* phage HP3

Viral genomes are highly diverse, and the understanding of viral evolutionary mechanisms is challenging (28). The diversity of phages is sometimes driven by PVPs (11). In this subsection, we used DeePVP to perform PVP annotation on *Escherichia* phage HP3 (RefSeq accession: NC_041920.1), a phage that may be an effective treatment strategy for murine models of bacteraemia (29), to reveal the genomic features of phage HP3. According to the annotation of the RefSeq database, the *Escherichia* phage HP3 genome encodes 269 proteins, all of which lack of the information of function annotation in RefSeq database. Additionally, no experimental data are available to determine which protein is PVP; therefore, PVP annotation using computational methods is an efficient way to analyse the genome. We used all the related tools to perform PVP annotation on the phage HP3 genome. The base coordinates for the predicted PVPs of each tool are shown in Figure 3. We defined a “PVP prediction reliability index” for each predicted PVP of each tool to evaluate the prediction reliability. For a certain protein predicted as PVP by a certain tool, if this protein was also predicted as PVP by the other *n* tools, then the “reliability index” for this predicted PVP was *n*. A higher *n* indicates that this protein is predicted as PVP by more tools at the same time, and therefore, the prediction may be more reliable. The average “reliability index” of each tool is shown in Figure 4, and we found that DeePVP achieved the highest index of 4.75, indicating that DeePVP might produce the most reliable predictions.

**Figure 3.**
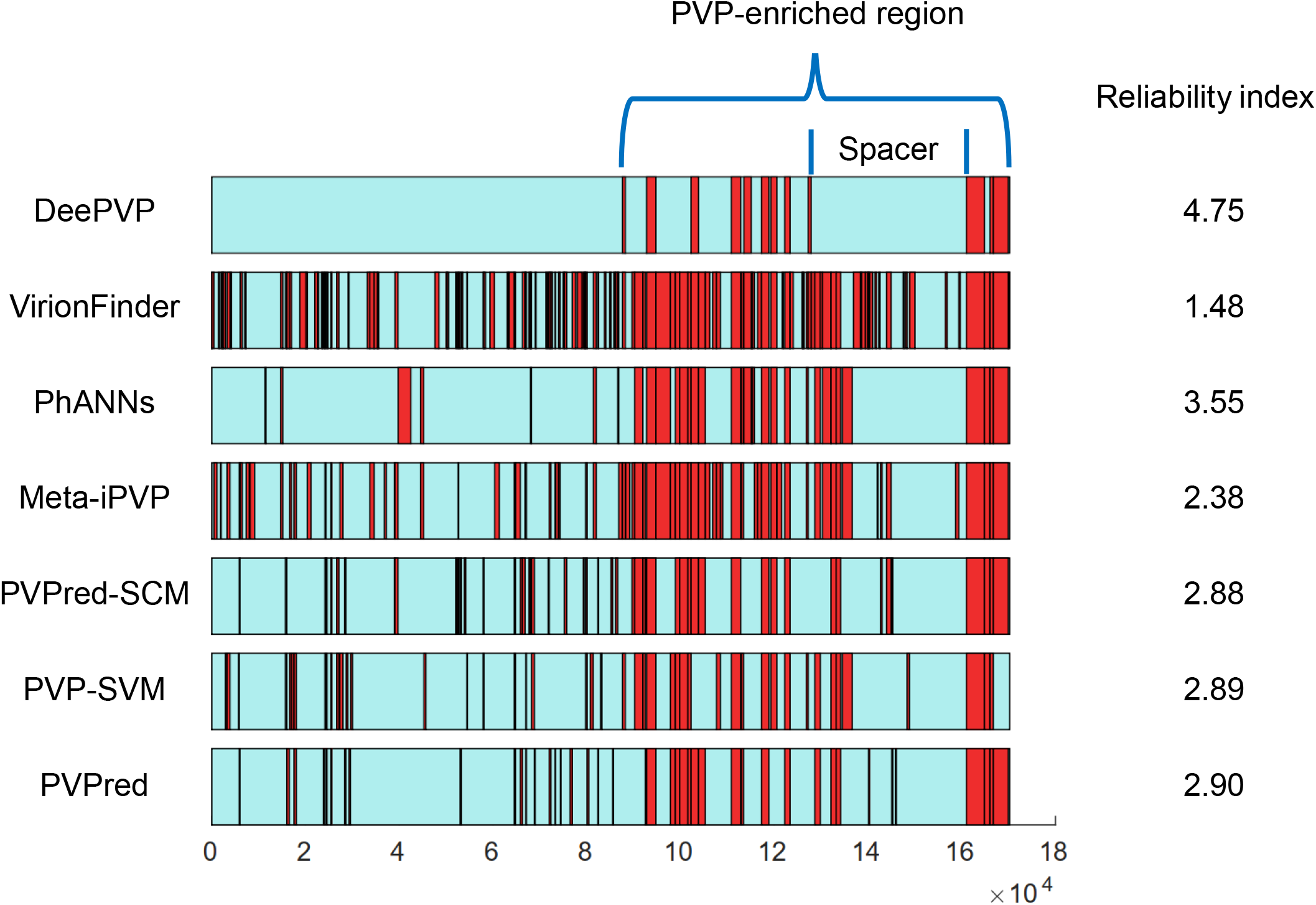
The distribution of PVPs predicted by each tool in the *Escherichia* phage HP3 genome. The blue bars represent the *Escherichia* phage HP3 genome, and the red boxes represent the predicted PVPs. The average reliability index of each tool is shown on the right side of the bar. The genomic features revealed by DeePVP are also marked in the first bar. The horizontal axis represents the base coordinates.

**Figure 4.**
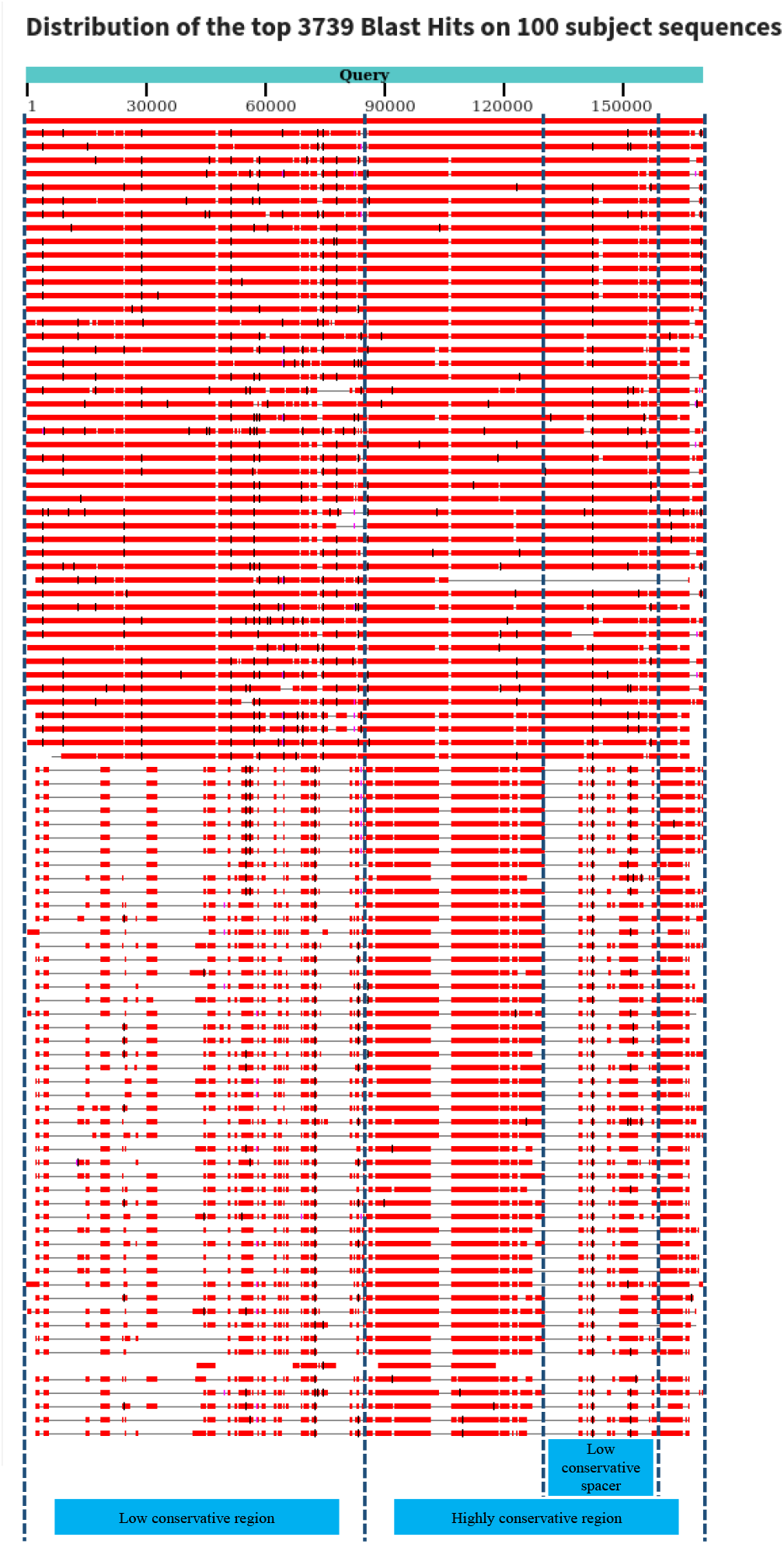
The alignment results of the top 100 target sequences for the *Escherichia* phage HP3 genome. The blue bar represents the query sequence of the phage HP3 genome. Each line below the query sequence represents a certain target sequence, and the coloured box on the target sequence represents a certain blast hit.

Additionally, we found that the genomic features of the phage HP3 genome revealed by DeePVP were interesting. Compared with the other tools, the distribution of all PVPs predicted by DeePVP was highly compact, and a PVP-enriched region was clearly present. Outside the PVP-enriched region, there were no predicted PVPs. This phenomenon is quite consistent with that of mycobacteriophage PDRPxv in application case 1, in which all PVPs are compactly encoded on the genome. Since a compact PVP distribution pattern is commonly observed (26,27), we consider that DeePVP may better reveal the genomic features of the phage genome. Meanwhile, from the DeePVP prediction results, we found that there was a clear spacer without predicted PVPs within the PVP-enriched region.

To further perform a comparative genomic analysis, we aligned the phage HP3 genome to the viral genomes in the NT database using the NCBI blastn online server (https://blast.ncbi.nlm.nih.gov/Blast.cgi), and all parameters were run under the default setting. The alignment results of the top 100 target sequences are shown in Figure 4, and detailed information on each target sequence is shown in Supplementary Data S2. The alignment graph showed that the phage HP3 genome contained a low conservative region that was conserved in some phage genomes but lacked obvious homology in other genomes and that it contained a highly conservative region that was present in almost all target sequences. In addition, within the highly conserved region, there was a low conservative spacer that was less conserved among these target sequences. More interestingly, when comparing the DeePVP prediction in Figure 3 with the alignment result in Figure 4, we found that the highly conservative region corresponded to the PVP-enriched region, while the low conservative region corresponded to the outside region of the PVP-enriched region. Additionally, the low conservative spacer within the highly conservative region approximately corresponded to the spacer within the PVP-enriched region. In addition to *Escherichia* phages, the target sequences also contained phages that infect other bacterial hosts, such as *Shigella, Enterobacteria, Serratia, Citrobacter, Yersinia*, and *Salmonella*. The above phenomenon suggests that the *Escherichia* phage HP3 genome contains a compact PVP-enriched region, which is conserved during viral evolution, and this PVP-enriched region contains a non-PVP spacer that might have been generated through recombination or horizontal gene transfer.

The PVP classification results of DeePVP and PhANNs are shown in Table 5 and Supplementary Data S3, respectively. We found that the predicted category of DeePVP could cover most of the essential PVP for the viral particle, including the major capsid, major tail, tail sheath, tail fiber, baseplate and portal. Additionally, three tail-associated PVPs, tail fibers, are encoded next to each other (YP_009965877.1, YP_009965879.1, YP_009965880.1). This phenomenon is similar to that of application case 1, in which five putative tail-associated PVPs, minor tails, are encoded continuously in the genome.

**Table 5.** PVP category predicted by DeePVP on the *Escherichia* phage HP3 genome.

| Protein ID | DeePVP prediction |
| --- | --- |
| YP_009965784.1 | Tail sheath |
| YP_009965791.1 | Baseplate |
| YP_009965797.1 | Tail fiber |
| YP_009965805.1 | Tail sheath |
| YP_009965807.1 | Portal |
| YP_009965812.1 | Major capsid |
| YP_009965814.1 | Major capsid |
| YP_009965818.1 | Major tail |
| YP_009965826.1 | Baseplate |
| YP_009965877.1 | Tail fiber |
| YP_009965879.1 | Tail fiber |
| YP_009965880.1 | Tail fiber |

## DISCUSSION

In this work, we present DeePVP to perform PVP annotation on a phage genome. The main module of DeePVP aims to identify whether a phage protein is PVP, while the extended module can further judge which class the PVP belongs to. Evaluation using the large-scale benchmark dataset shows that DeePVP performs much better than the related tools. We provided two application cases to demonstrate the value of DeePVP. In the case of mycobacteriophage PDRPxv, by referring to the experimental data of mass spectrometry, we illustrated that the prediction of DeePVP was more reliable and could better reveal the compact distribution pattern of PVP over the genome. We then used DeePVP to perform PVP prediction on *Escherichia* phage HP3, which lacked experimental data and annotation information for its PVPs. Compared with the other tools, DeePVP again showed a clear PVP-enriched region over the genome, and we found that this newly discovered PVP-enriched region may be conserved during the viral evolution process. We therefore suggest that DeePVP may be a powerful tool for phage genomic analysis and understanding the viral evolution process.

In DeePVP, we chose to train a two-class classifier and a ten-class classifier separately rather than train an eleven-class classifier directly. This is because the number of protein sequences in each category of the dataset is unbalanced. For example, in the dataset, 369,553 of the proteins belong to the non-PVP category, while only 2,105 proteins belong to the collar category. An unbalanced dataset is challenging for training the neural network because the neural network may tend to judge most of the samples to the category with the largest size automatically. In DeePVP, we first trained a CNN to separate PVP and non-PVP; therefore, we can reduce the impact of the non-PVP category, which contains many more sequences than the other PVP categories, on the PVP classification task.

In the related tools, amino acid composition-based feature vectors are a commonly used method to characterize sequences. Such global statistics may lose some local information, such as some conserved domains or motifs, of the sequence. Additionally, such feature vector may be sparse. For example, PhANNs uses a *k*-mer-based feature vector containing thousands of bits, while phage proteins are often shorter than that of bacterial (9) and a large number of proteins contain only approximately 250 aa; therefore, most of the bits in the feature vector may be 0. Such sparse vectors may make it difficult for the algorithm to fit the data. In DeePVP, we used the “one-hot” form to encode the sequence, and the information of each amino acid remained in the characterization model. We used CNN to extract the useful features from the raw data. It has been shown that CNN is powerful for extracting useful features, and the convolution kernel may serve as the sensitive position weight matrix (PWM) to detect local specific motifs (30-32). We therefore consider that such a method may help to improve the performance of DeePVP.

Although the benchmark dataset we used in this work is the largest and well-designed PVP and non-PVP dataset by far, such a dataset is primarily focused on the 10 major classes of PVP and was created using keyword search on the public database. Thus, some less frequently occurring PVP may be excluded in the dataset, leading to DeePVP failing to identify some kinds of PVP. For example, in application case 1, DeePVP failed to identify a putative scaffolding protein (Gp11) and a putative tape measure protein (Gp28), which may be because these two kinds of proteins were not included in the dataset during the keyword search process. Therefore, more effort should be made to create more exhaustive PVP sets to further improve the algorithm performance.

Although the comprehensive index *F1-score* of DeePVP is much better than that of the other tools, the *recall* of DeePVP is slightly lower than that of some tools, such as PhANNs and VirionFinder, in the PVP identification task, indicating that DeePVP may fail to identify some PVPs. On the other hand, DeePVP had the better ability to reveal the PVP-enriched region, and PVPs in many phages are encoded compactly within the region. Therefore, to identify as much PVP as possible under a low false positive prediction rate, we suggest that users can combine the usage of different tools. In application case 1, for example, the user can first use DeePVP to identify the PVP-enriched region (from Gp8 to Gp33, as shown in Figure 2 and Table 4). Within the region, users can further use the more sensitive tool PhANNs to identify more PVPs. Since proteins within the region are more likely to be PVP, such an operation may not generate much false positive predictions. In this way, the PVP of Gp28, which fails to be predicted by DeePVP, will be included in the prediction, and the *recall* will increase from 75% to 83.33%. Although such an operation will also introduce three false positive predictions (Gp14, Gp19, Gp26), a large number of false positive predictions outside the PVP-enriched region are excluded. In the future, it is worth constructing a comprehensive workflow integrating different tools to improve the PVP annotation performance.

DeePVP is primarily designed for PVP annotation on isolated phage genomes. Recently, a large amount of metagenomic data containing both bacterial DNA and viral DNA has emerged. Such data also contain incomplete proteins because of poor sequence assembly. More effort should also be made to allow the algorithm to identify PVP from a large number of bacterial proteins in future work.

## DATA AVAILABILITY

The training and testing data of DeePVP can be downloaded from http://edwards.sdsu.edu/phanns/download/expandedDB.tgz. The mycobacteriophage PDRPxv and *Escherichia* phage HP3 genome can be downloaded from NCBI using the accession of KR029087 and NC_041920.1, respectively. DeePVP is freely available at https://github.com/zhenchengfang/DeePVP/.

## CONFILICT OF INTEREST

The authors declare no competing interests.

