## Supplementary Data S1 for "DeePVP: Identification and classification of phage virion protein using deep learning"

**1 “One-hot” encoding form**

In DeePVP, each amino acid is represented by a “one-hot” vector with 20 bits, as mentioned below:

A:	[0 0 0 0 0 0 0 0 0 0 0 0 0 0 0 0 0 0 0 1]

C:	[0 0 0 0 0 0 0 0 0 0 0 0 0 0 0 0 0 0 1 0]

D:	[0 0 0 0 0 0 0 0 0 0 0 0 0 0 0 0 0 1 0 0]

E:	[0 0 0 0 0 0 0 0 0 0 0 0 0 0 0 0 1 0 0 0]

F:	[0 0 0 0 0 0 0 0 0 0 0 0 0 0 0 1 0 0 0 0]

G:	[0 0 0 0 0 0 0 0 0 0 0 0 0 0 1 0 0 0 0 0]

H:	[0 0 0 0 0 0 0 0 0 0 0 0 0 1 0 0 0 0 0 0]

I:	[0 0 0 0 0 0 0 0 0 0 0 0 1 0 0 0 0 0 0 0]

K:	[0 0 0 0 0 0 0 0 0 0 0 1 0 0 0 0 0 0 0 0]

L:	[0 0 0 0 0 0 0 0 0 0 1 0 0 0 0 0 0 0 0 0]

M:	[0 0 0 0 0 0 0 0 0 1 0 0 0 0 0 0 0 0 0 0]

N:	[0 0 0 0 0 0 0 0 1 0 0 0 0 0 0 0 0 0 0 0]

P:	[0 0 0 0 0 0 0 1 0 0 0 0 0 0 0 0 0 0 0 0]

Q:	[0 0 0 0 0 0 1 0 0 0 0 0 0 0 0 0 0 0 0 0]

R:	[0 0 0 0 0 1 0 0 0 0 0 0 0 0 0 0 0 0 0 0]

S:	[0 0 0 0 1 0 0 0 0 0 0 0 0 0 0 0 0 0 0 0]

T:	[0 0 0 1 0 0 0 0 0 0 0 0 0 0 0 0 0 0 0 0]

V:	[0 0 1 0 0 0 0 0 0 0 0 0 0 0 0 0 0 0 0 0]

W:	[0 1 0 0 0 0 0 0 0 0 0 0 0 0 0 0 0 0 0 0]

Y:	[1 0 0 0 0 0 0 0 0 0 0 0 0 0 0 0 0 0 0 0]

For a given protein, the “one-hot” vector of each amino acid will be joined together according to the amino acid sequence order. Therefore, a protein with length *L* can be represented by a “one-hot” matrix of length *L* and width 20. The convolutional neural network needs to fix the size of the input matrix. We found that 99.83% of the protein in the benchmark dataset was shorter than 2000 aa. To save computational resources, we filter out the amino acids downstream of the 2000^th^ aa. For sequences shorter than 2000 aa, the “one-hot” matrix will be padded with zero vectors to 2000 rows.

**2 Hyperparameter selection for the convolutional neural network**

**2.1 The mail module**

Hyperparameters in the 1D convolution layer: the length of the convolution layer was set to 9, the number of convolutional kernels was set to 700, and the “ReLU” activation function was used.

Hyperparameters in the full connection layer: the number of nodes was set to 700, and the “ReLU” activation function was used.

Other hyperparameters: In the training process, two dropout layers were deposited between the batch normalization layer and the full connection layer and between the full connection layer and the sigmoid layer. Both dropout probability was set to 0.55. The loss function of “binary cross-entropy” was used. The optimizer of “Adam” was used. The number of epochs was set to 80, and the batch size was set to 200.

**2.2 The extended module**

Hyperparameters in the 1D convolution layer: the length of the convolution layer was set to 8, the number of convolutional kernels was set to 600, and the “ReLU” activation function was used.

Hyperparameters in the full connection layer: the number of nodes was set to 600, and the “ReLU” activation function was used.

Other hyperparameters: In the training process, two dropout layers were deposited between the batch normalization layer and the full connection layer and between the full connection layer and the softmax layer. Both dropout probability was set to 0.5. The loss function of “categorical cross-entropy” was used. The optimizer of “Adam” was used. The number of epochs was set to 80, and the batch size was set to 256.
